# Yeast Nuak1 phosphorylates histone H3 threonine 11 in low glucose stress conditions by the cooperation of AMPK and CK2 signaling

**DOI:** 10.1101/2020.11.03.367094

**Authors:** Seunghee Oh, Jaehyoun Lee, Selene K. Swanson, Laurence Florens, Michael P. Washburn, Jerry L. Workman

## Abstract

Changes in available nutrients are inevitable events for most living organisms. Upon nutritional stress, several signaling pathways cooperate to change the transcription program through chromatin regulation to rewire cellular metabolism. In budding yeast, histone H3 threonine 11 phosphorylation (H3pT11) acts as a marker of low glucose stress and regulates the transcription of nutritional stress responsive genes. Understanding how this histone modification ‘senses’ external glucose changes remains elusive. Here, we show that Tda1, the yeast orthologue of human Nuak1, is a direct kinase for H3pT11 upon low glucose stress. Yeast AMPK directly phosphorylates Tda1 to govern Tda1 activity, while CK2 regulates Tda1 nuclear localization. Collectively, AMPK and CK2 signaling converge on histone kinase Tda1 to link external low glucose stress to chromatin regulation.

## Introduction

To ensure survival, cells must properly adapt to changes in available nutrients by altering their metabolism. This adaptation can be achieved through the cooperation of multiple metabolic pathways. Among those pathways, AMPK (AMP-activated protein kinase) signaling has a central role in energy homeostasis, especially when the cellular nutrient supply is low (González, Hall, Lin, & Hardie, 2020). AMPK orthologues are very well-conserved from yeast to human, and occur as heterotrimeric complex comprising catalytic α subunit and regulatory β and γ subunits (Ross, MacKintosh, & Hardie, 2016). For its proper function, the AMPK catalytic α subunit needs to be phosphorylated at its conserved threonine residue (threonine 172 in rat (Hawley et al., 1996), and threonine 210 in budding yeast (McCartney & Schmidt, 2001)) within the activation loop of its kinase domain. In budding yeast, the catalytic α subunit is Snf1 (Sucrose Non-Fermenting 1). This name comes from its requirement for growth by sucrose fermentation (Carlson, Osmond, & Botstein, 1981). The regulatory γ subunit of budding yeast AMPK is Snf4, and its interaction with Snf1 liberates Snf1 from auto-inhibition that interferes with Snf1 threonine 210 (T210) phosphorylation (Chen et al., 2009). Budding yeast encodes three AMPK β subunit proteins; Sip1, Sip2 and Gal83. They are partially redundant. Only when all three β subunits are deleted, cells exhibit growth defects in media containing ethanol or glycerol as a sole carbon source, as the *snf1Δ* mutant does (Erickson & Johnston, 1993; Schmidt & McCartney, 2000). The β subunits have a conserved C terminal sequence that interacts with α and γ subunits (Jiang & Carlson, 1997; X. Yang, Jiang, & Carlson, 1994) and a specific N terminal sequence for each β subunit which confers a distinctive subcellular localization of the Snf1 complex. Notably, Gal83 is required for Snf1 nuclear localization upon glucose depletion (Olivier Vincent, Townley, Kuchin, & Carlson, 2001).

Snf1 interacts with and phosphorylates several proteins involved in nutritional stress responses (Coccetti, Nicastro, & Tripodi, 2018). Its substrates include several transcription factors such as Cat8 and Sip4 which bind to Carbon Source Responsive Elements (CSRE) for the transcription of gluconeogenic genes (Lesage, Yang, & Carlson, 1996; Randez-Gil, Bojunga, Proft, & Entian, 1997; O. Vincent & Carlson, 1998). Cat8 is required for proper expression of the transcription factor Adr1, which is also phosphorylated by Snf1 (Kacherovsky, Tachibana, Amos, Fox, & Young, 2008; Young, Kacherovsky, & Van Riper, 2002). Snf1 also phosphorylates the Mig1 transcription repressor, which suppresses the transcription of glucose repressive genes (Nehlin & Ronne, 1990). Mig1 phosphorylation by Snf1 activates its nuclear export signal leading to expulsion of Mig1 from the nucleus (DeVit & Johnston, 1999; Papamichos-Chronakis, Gligoris, & Tzamarias, 2004; Treitel, Kuchin, & Carlson, 1998). Interestingly, the yeast hexokinase 2 (Hxk2) protein interacts with Mig1 in the nucleus to protect Mig1 from phosphorylation by Snf1 (Ahuatzi, Riera, Pelaez, Herrero, & Moreno, 2007). When glucose becomes scarce, Hxk2 is phosphorylated at serine 15 which inactivates its nuclear localization signal and, in turn, facilitates its cytoplasmic localization (Fernandez-Garcia, Pelaez, Herrero, & Moreno, 2012; Kriegel, Rush, Vojtek, Clifton, & Fraenkel, 1994). Snf1 had been believed to be a Hxk2 serine 15 kinase, however, recent studies revealed that yeast Nuak1 homolog Tda1 is responsible for Hxk2 phosphorylation (Kaps et al., 2015; Kettner et al., 2012). Thus far, Hxk2 is the only known substrate of Tda1, and how Tda1 activity is regulated upon low glucose stress is unknown.

AMPK also has a central role in regulating nutritional stress-specific chromatin modification (Lee, Oh, Abmayr, & Workman, 2020). In yeast, Snf1 phosphorylates H3 at serine 10 at the *INO1* gene promoter to regulate its transcription (Lo et al., 2001). Human AMPK phosphorylates H2B at serine 36 at the promoters of p53 responsive genes upon glucose starvation (Bungard et al., 2010). AMPK indirectly regulates H3 arginine 17 di-methylation by affecting histone arginine methyltransferase CARM1 protein level in the nucleus upon apoptosis-inducing conditions (Shin et al., 2016).

Recent studies imply that metabolic enzymes themselves also participate in chromatin regulation. Pyruvate kinase, a glycolysis enzyme, can phosphorylate H3 at threonine 11 in humans and in yeast (Li et al., 2015; W. Yang et al., 2012). Especially in yeast, pyruvate kinase forms a complex named SESAME with several metabolic enzymes involved in serine and SAM metabolism to regulate H3 threonine 11 phosphorylation (H3pT11) in glucose rich conditions (Li et al., 2015). Interestingly, we found that H3pT11 can act as a marker of low glucose stress in yeast (Oh, Suganuma, Gogol, & Workman, 2018). The global H3pT11 level is inversely correlated with external glucose level and specifically increased at genes involved in metabolic changes. Notably, the H3pT11 increase upon low glucose stress is a SESAME-independent process, indicating multiple different kinases are involved in H3pT11 regulation. We found that Cka1, a catalytic subunit of CK2 complex, is required for H3pT11 regulation in low glucose conditions. However, CK2 has been observed to be constitutively active and insensitive to external changes of environmental cues (Pinna, 2002). It remains elusive how a constantly active CK2 can regulate H3pT11, which becomes elevated upon low glucose stress. This suggests that H3pT11 may require additional mechanisms for sensing the availability of carbon sources.

In this study, we identify an understudied kinase, Tda1, as a histone H3 T11 kinase in low glucose stress conditions. Snf1 directly phosphorylates Tda1 at its C-terminus, especially at tandem serine 483 and threonine 484 (S483/T484) residues, which is required for *in vivo* Tda1 activity. CK2 regulates H3pT11 by controlling Tda1 nuclear localization. Hence, Tda1 acts as a signaling platform that can combine Snf1 and CK2 signaling, thereby connecting external nutrient availability to transcription regulation in the nucleus.

## Results

### H3pT11 phosphorylation upon low glucose stress is dependent on Snf1

We previously reported a genome-wide study that histone H3 threonine 11 phosphorylation (H3pT11) specifically increases at the promoters of stress responsive genes upon low glucose stress, and H3pT11 is required for proper transcription of those genes (Oh et al., 2018). As AMPK signaling has major regulatory roles in nutrient starvation conditions (González et al., 2020), we tested whether H3pT11 is linked with the yeast AMPK homolog, Snf1 signaling pathway. Upon the media shift from glucose rich (2%) YPD to nutritionally unfavorable YP with 3% glycerol (YPgly), Snf1 threonine 210 phosphorylation (Snf1 pT210), the active Snf1 marker, showed a similar pattern of increase compared to that of H3pT11 (Figure 1A). In addition, a *snf1Δ* mutant showed significantly reduced global H3pT11 levels in saturated cultures (Figure 1B) where media glucose is depleted (Oh et al., 2018), and impaired H3pT11 increase upon media shift from YPD to YPgly (Figure 1C). Rim15 is another kinase which becomes active in low glucose conditions (Vidan & Mitchell, 1997; Wei et al., 2008), but a *rim15Δ* mutant did not show the global H3pT11 defect (Figure 1B). Snf1 phosphorylates several transcription factors including Adr1, Cat8, Sip4 and Mig1 upon nutritional stress (Kacherovsky et al., 2008; Lesage et al., 1996; Randez-Gil et al., 1997; Treitel et al., 1998), however, deletion of these transcription factors did not affect H3pT11 levels (Figure 1-figure supplement 1). These results suggest that H3pT11 is specifically regulated via the Snf1 pathway and reduction of H3pT11 in a *snf1Δ* mutant is not an indirect result of a transcriptional defect. Previously, we showed that Cka1, the catalytic subunit of CK2, is required for the H3pT11 increase in low glucose stress conditions (Oh et al., 2018). Interestingly, a *cka1Δsnf1Δ* double deletion mutant showed a similar level of H3pT11 compared to a *snf1Δ or cka1Δ* single deletion mutant (Figure 1D), suggesting that CK2 and Snf1 may act in the same pathway for H3pT11 regulation.

**Figure 1.**
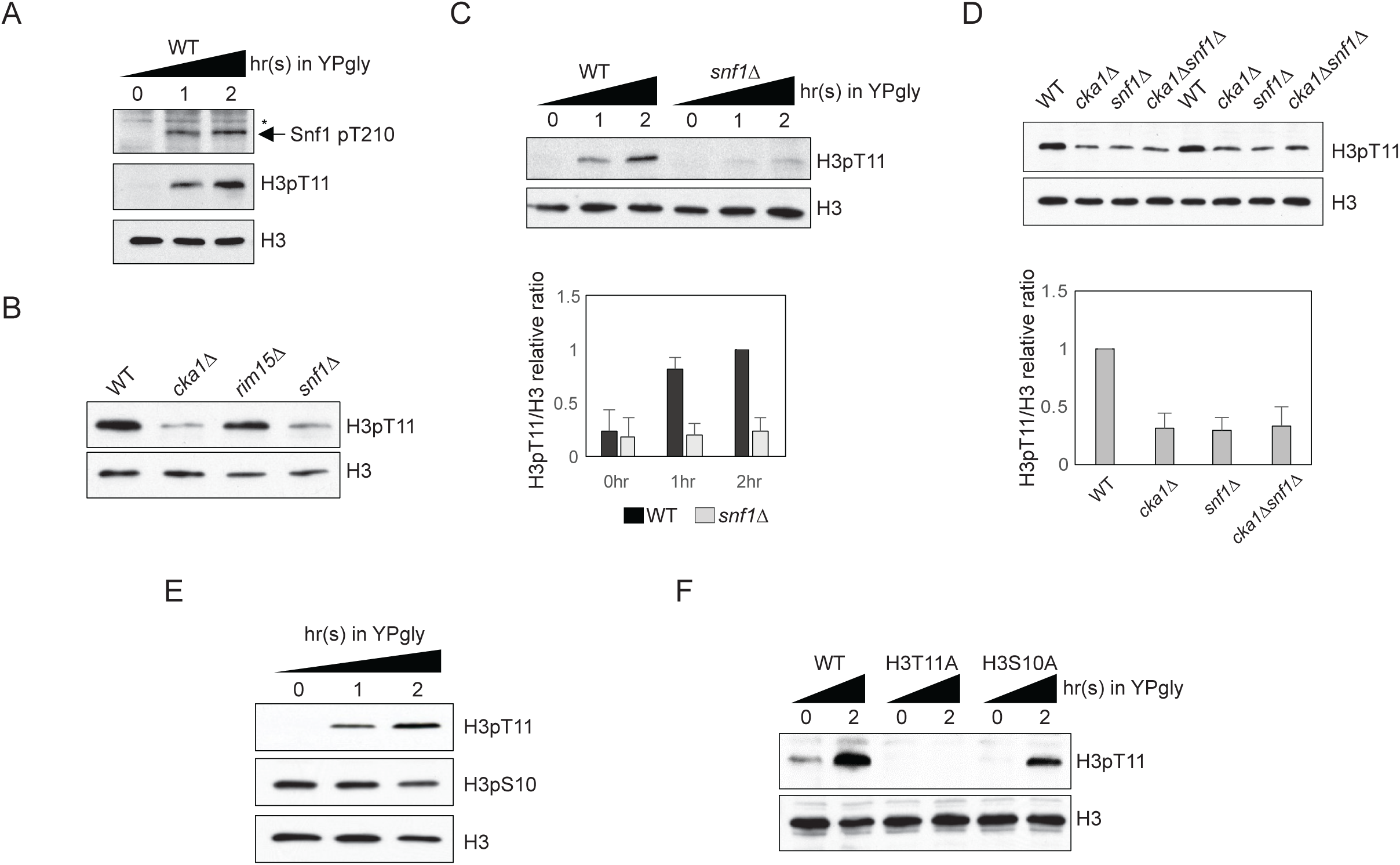
H3pT11 is an AMPK dependent, but H3pS10 independent histone modification upon low glucose stress. (A) Comparison of the Snf1 threonine 210 phosphorylation (Snf1 pT210) and H3pT11 levels upon media shift from YPD (2% glucose) to YPgly (YP with 3% glycerol) measured by western blots. (B) Global H3pT11 levels in WT (BY4741), *cka1Δ, rim15Δ*, and *snf1Δ* cells measured by western blot. The cells were taken from overnight saturated cultures in YPD media. (C) (Upper panel) Comparison of H3pT11 levels in WT and *snf1Δ* upon the media shift from YPD to YPgly at indicated time points analyzed by western blots. (Lower panel) Relative ratios of H3pT11 to H3 signals presented with error bars indicating standard deviations (STD) of three biological replicates. (D) (Upper panel) Global H3pT11 levels in WT, *cka1Δ, snf1Δ*, and *cka1Δsnf1Δ* cells taken from saturated cultures in YPD media measured by western blots. (Lower panel) Relative band intensities of H3pT11 to H3 signals. Error bars indicate STD from three biological replicates. (E) Changes in H3pT11 and H3pS10 signals in the WT strain (BY4741) upon the media shift from YPD to YPgly at indicated time points measured by western blots. (F) Comparison of H3pT11 upon media shift from YPD to YPgly in WT (y1166), H3T11A, and H3S10A strains analyzed by western blots.

Snf1 can phosphorylate histone H3 at serine 10 in yeast (Lo et al., 2001). As H3 serine 10 is next to H3 T11, we tested whether histone H3 S10 phosphorylation (H3pS10) and H3pT11 behave similarly. Upon media shift from YPD to YPgly, the global H3pS10 level remained stable, while H3pT11 was increased (Figure 1E). In addition, an H3S10A mutant, where histone H3 serine 10 is mutated into alanine, showed an unperturbed H3pT11 increase upon glucose depletion (Figure 1F). These results indicate that H3pT11 and H3pS10 are differently regulated under low glucose stress, and Snf1 signaling, but not H3pS10, is required for H3pT11 increase in the stress condition.

### Snf1 and CK2 are not direct kinases for H3pT11

Snf1 forms a functional AMPK complex with the regulatory β and γ subunits (Ross et al., 2016). When we measured H3pT11 levels from Snf1 complex subunit deletion mutants, interestingly, only the catalytic α subunit mutant, *snf1Δ*, showed a defect while β (Gal83, Sip1, and Sip2) and γ subunit (Snf4) deletion mutants showed a comparable level of H3pT11 to WT (Figure 2A). Indeed, a β subunit mutant (*gal83Δ*) showed a slightly increased level of H3pT11. As Gal83 dictates Snf1 nuclear localization upon low glucose stress (Olivier Vincent et al., 2001), this result raised the possibility that Snf1 may not be a direct kinase or a major kinase for H3pT11 *in vivo*.

**Figure 2.**
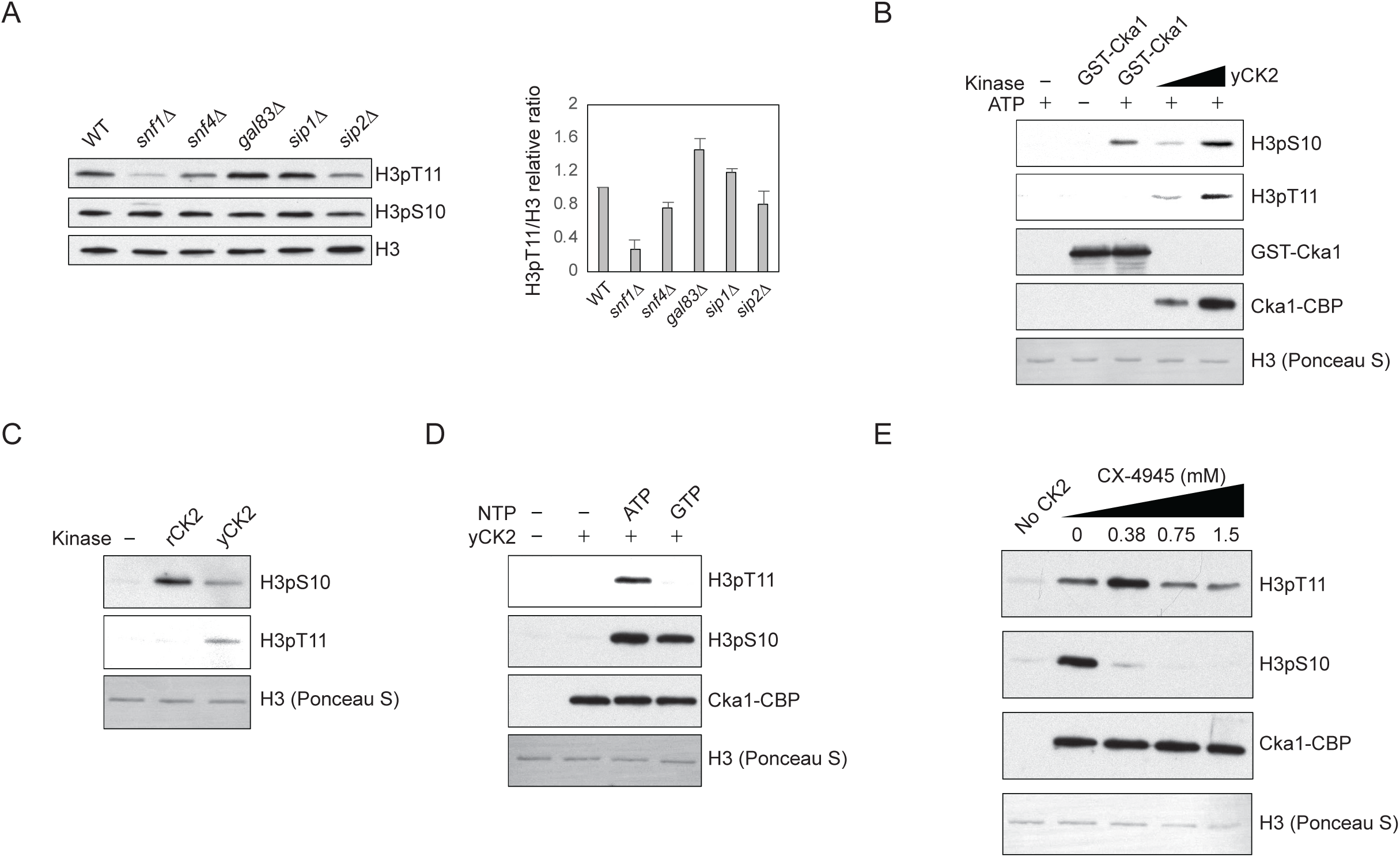
Snf1 and CK2 are not direct kinases for H3pT11. (A) (Left) Global H3pT11 and H3pS10 levels of Snf1 complex α subunit (Snf1), γ subunit (Snf4), and β subunit (Gal83, Sip1, and Sip2) deletion mutants compared to WT (BY4741) analyzed by western blots. Cells were taken from saturated cultures in YPD media. (Right) The relative band intensities of H3pT11 to H3 with error bars indicating STD of three biological replicates. (B) *In vitro* kinase assay of recombinant GST-Cka1 and yeast TAP purified CK2 (yCK2) using recombinant Xenopus histone H3 as a substrate. (C) *In vitro* kinase assay of human recombinant CK2 complex (rCK2) and yCK2 using recombinant H3 as a substrate. (D) *In vitro* kinase assay of yCK2 with recombinant H3 as a substrate and ATP or GTP as a phosphate donor. (E) *In vitro* kinase assay of yCK2 for H3 with increasing amount of CX-4945 treatment.

Our previous study showed that yeast TAP purified CK2 can phosphorylate H3 at T11 *in vitro* (Oh et al., 2018). However, recombinant yeast Cka1 and recombinant human CK2 complex did not phosphorylate H3 at T11 *in vitro*, while both TAP purified and recombinant CK2 phosphorylated H3 at S10 (Figures 2B and 2C). CK2 is unusual for a kinase in that it can utilize both ATP and GTP as phosphate donors (Niefind, Pütter, Guerra, Issinger, & Schomburg, 1999). When GTP was used as the phosphate donor, TAP purified CK2 phosphorylated H3 at S10, but not at T11 (Figure 2D). In addition, the CK2 specific inhibitor CX-4945 (Siddiqui-Jain et al., 2010) suppressed CK2 activity against H3pS10, but not against H3pT11 *in vitro* (Figure 2E). These results indicate that an unknown CK2 interacting kinase, rather than CK2 itself, is responsible for H3pT11 *in vitro*.

### Tda1 is responsible for H3pT11 upon low glucose stress *in vivo*

Since Snf1 and CK2 are not direct kinases against H3pT11, we inquired as to which kinase might be responsible for the modification. As the Snf1 pathway is critical in nutritional stress conditions, we hypothesized that Snf1 interacting kinases may be responsible or participate in H3pT11 regulation. Using the Saccharomyces Genome Database (https://www.yeastgenome.org) as a guide, we tested H3pT11 levels of deletion mutants of the kinases which are known to interact with the Snf1 complex. Unexpectedly, the yeast Nuak1 homolog *tda1Δ* mutant showed a significantly reduced H3pT11 levels (Figure 3A). In glucose depleted media, the *tda1Δ* mutant showed a more severe defect in the global H3pT11 levels than the *snf1Δ* mutant (Figure 3B). H3pS10 levels did not significantly change in *tda1Δ*, supporting the independence of H3pT11 from H3pS10 in this stress condition. In media-shift assays, the *tda1Δ* mutant showed impaired H3pT11 increase in YPgly, similar to *snf1Δ* (Figure 3C). Interestingly, upon media shift from YPD to YPgly, Tda1 protein levels were significantly increased in both the cytoplasm and the nucleus in the wildtype strain (Figure 3D and Figure 3-figure supplement 1). Collectively, these results strongly suggest that Tda1 has a central role in H3pT11 regulation upon low glucose stress.

**Figure 3.**
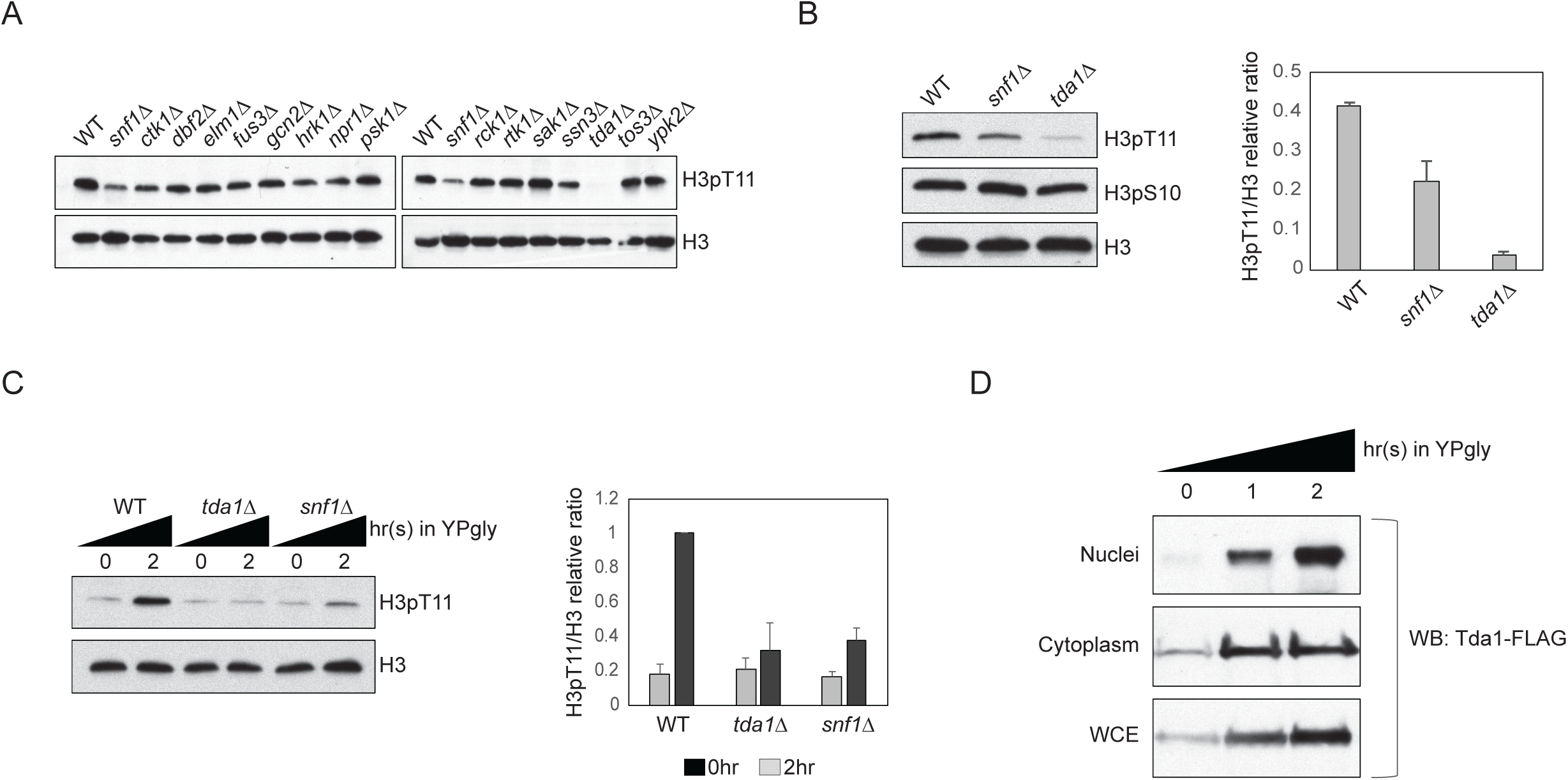
Tda1 is responsible for H3pT11 upon low glucose stress *in vivo*. (A) Global H3pT11 levels in Snf1-interacting kinase mutant cells taken from saturated cultures in YPD media measured by western blots. (B and C) (Left panels) H3pT11 levels in WT, *snf1Δ*, and *tda1Δ* cells taken from (B) saturated cultures in YPD media or taken at (C) indicated time points upon media shift from YPD to YPgly analyzed by western blots. (Right panels) The relative band intensities of H3pT11 to H3 are presented with error bars indicating STD of three biological replicates. (D) Tda1 protein levels tagged with C-terminal 3xFLAG tag in the nuclei, cytoplasm, and whole cell extracts (WCE) upon media shift from YPD to YPgly media at indicated time points measured by western blots against FLAG tag.

### Tda1 phosphorylates H3 at T11 *in vitro*

Thus far, the only known Tda1 target is yeast hexokinase2 (Hxk2) at serine 15 (Kaps et al., 2015; Kettner et al., 2012). We noticed that the surrounding sequence of Hxk2 serine 15 is similar to that of histone H3 T11 (Figure 4A). To test if Tda1 can directly phosphorylate H3 *in vitro*, we purified Tda1 from yeast using TAP tag purification (Figure 4B). TAP purified Tda1 did not show many bands in a PAGE gel, suggesting that Tda1 may not be a part of multi-subunit complex. An *in vitro* assay showed that indeed, TAP purified Tda1 protein can robustly phosphorylate H3 at T11 (Figure 4C). Recombinant GST tagged Tda1 purified from *E. coli* also phosphorylated H3 at T11 but not at S10, while Snf1 purified from yeast using a FLAG tag phosphorylated H3 at both S10 and T11 *in vitro* (Figure 4D). Sequence analysis from Saccharomyces Genome Database (http://www.yeastgenome.org) predicts that Tda1 may contain a kinase domain at the N-terminus. In this regard, we made several Tda1 fragments (Tda1 N1 – N5) to define the kinase domain (Figure 4E). An *in vitro* kinase assay of the fragments showed that Tda1 N1 (amino acids 1 to 353) did not phosphorylate H3 at T11 unlike Tda1 N2 (amino acids 1 to 380). Interestingly, Tda1 N3 (amino acids 1 to 426) showed a reduced activity against H3pT11 compared to Tda1 N2 (Figure 4F). These results suggest that Tda1 catalytic domain resides in amino acid 1 to 380, and the amino acids patch spanning 381 to 426 has a potential inhibitory effect on Tda1 activity against H3pT11.

**Figure 4.**
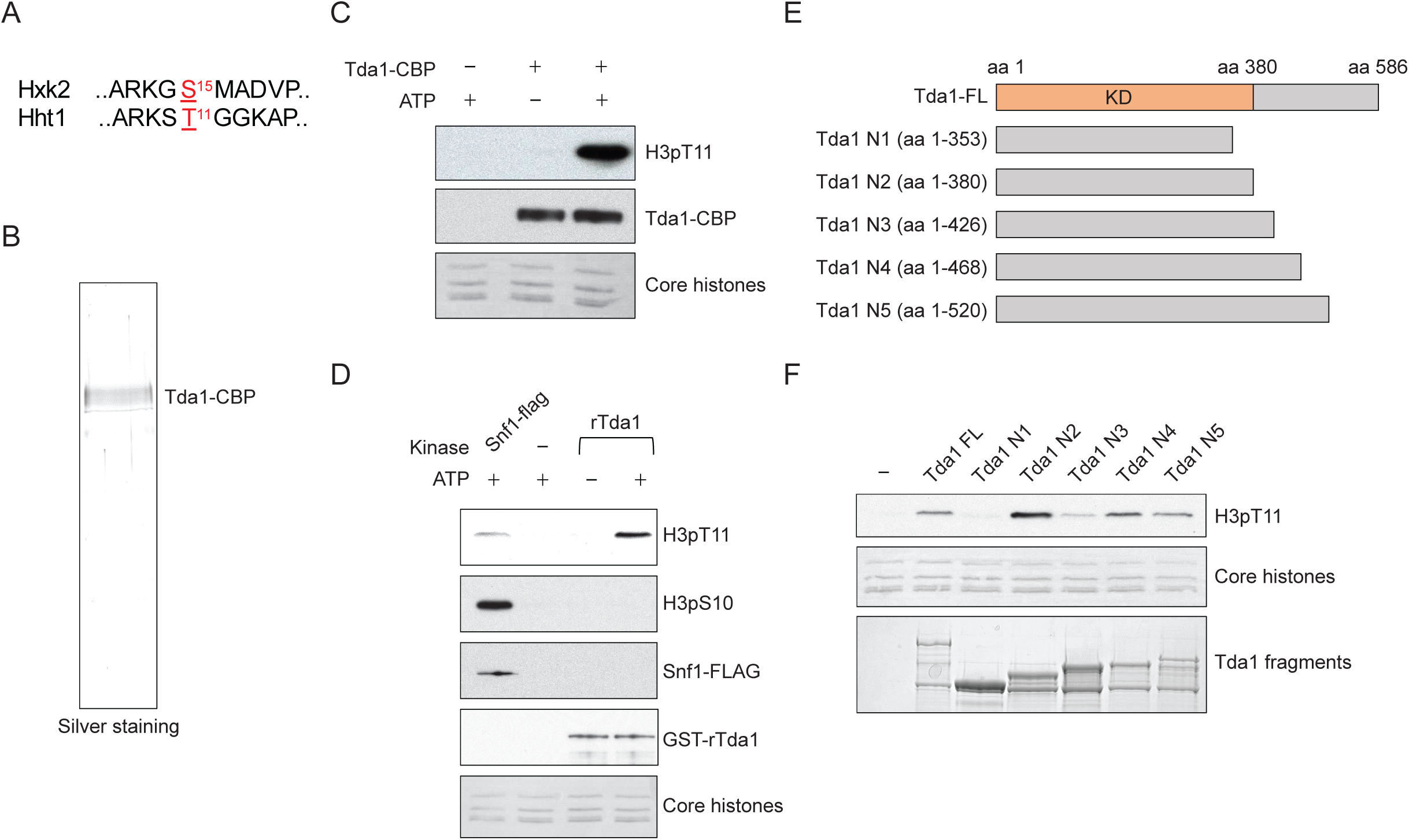
Tda1 phosphorylates H3 at T11 *in vitro*. (A) Comparison between the surrounding sequences of Hxk2 serine 15 and H3 (Hht1) threonine 11. (B) Silver staining of TAP purified Tda1 protein. (C) *In vitro* kinase assay of TAP purified Tda1 using core histones (H2A, H2B, H3, and H4) as substrates. (D) *In vitro* kinase assay of Snf1 complex purified from yeast using FLAG tag and recombinant yeast GST-Tda1 purified from *E*.*coli* using core histones (H2A, H2B, H3, and H4) as substrates. (E) Schematic diagram of recombinant GST-Tda1 N-terminal fragments used in (F). (F) *In vitro* kinase assay of Tda1 N fragments using core histones as substrates.

### Snf1 directly phosphorylates Tda1 at C-terminus

Next, we inquired how Snf1 signaling can regulate the activity of Tda1 towards H3pT11. irst, we compared the changes in Tda1 protein levels upon media shift from YPD to YPgly in WT and *snf1Δ* (Figure 5A). Interestingly, Tda1 showed multiple species migrating differently on a PAGE gel. In a low glucose stress condition, slowly migrating Tda1 species became dominant in the WT strain. However, in the *snf1Δ* mutant, the faster migrating species was dominant. In the *cka1Δ* mutant, Tda1 protein level changes were similar to that of WT (Figure 5-figure supplement 1A). We speculated that the apparent size differences of Tda1 in WT and *snf1Δ* reflected differential post translational modification patterns of Tda1. To test this possibility, Tda1 was purified from yeast using FLAG tag, then incubated with λ phosphatase. After the phosphatase treatment, Tda1 showed a more distinctive one band pattern while the slower migrating bands disappeared, indicating Tda1 is indeed a phosphorylated protein (Figure 5B). In addition, when we compared Tda1 purified from WT, *snf1Δ*, and *cka1Δ* backgrounds, the phosphorylated species of Tda1 was significantly reduced specifically in the *snf1Δ* mutant, implying that Snf1 is responsible for Tda1 phosphorylation (Figure 5C).

**Figure 5.**
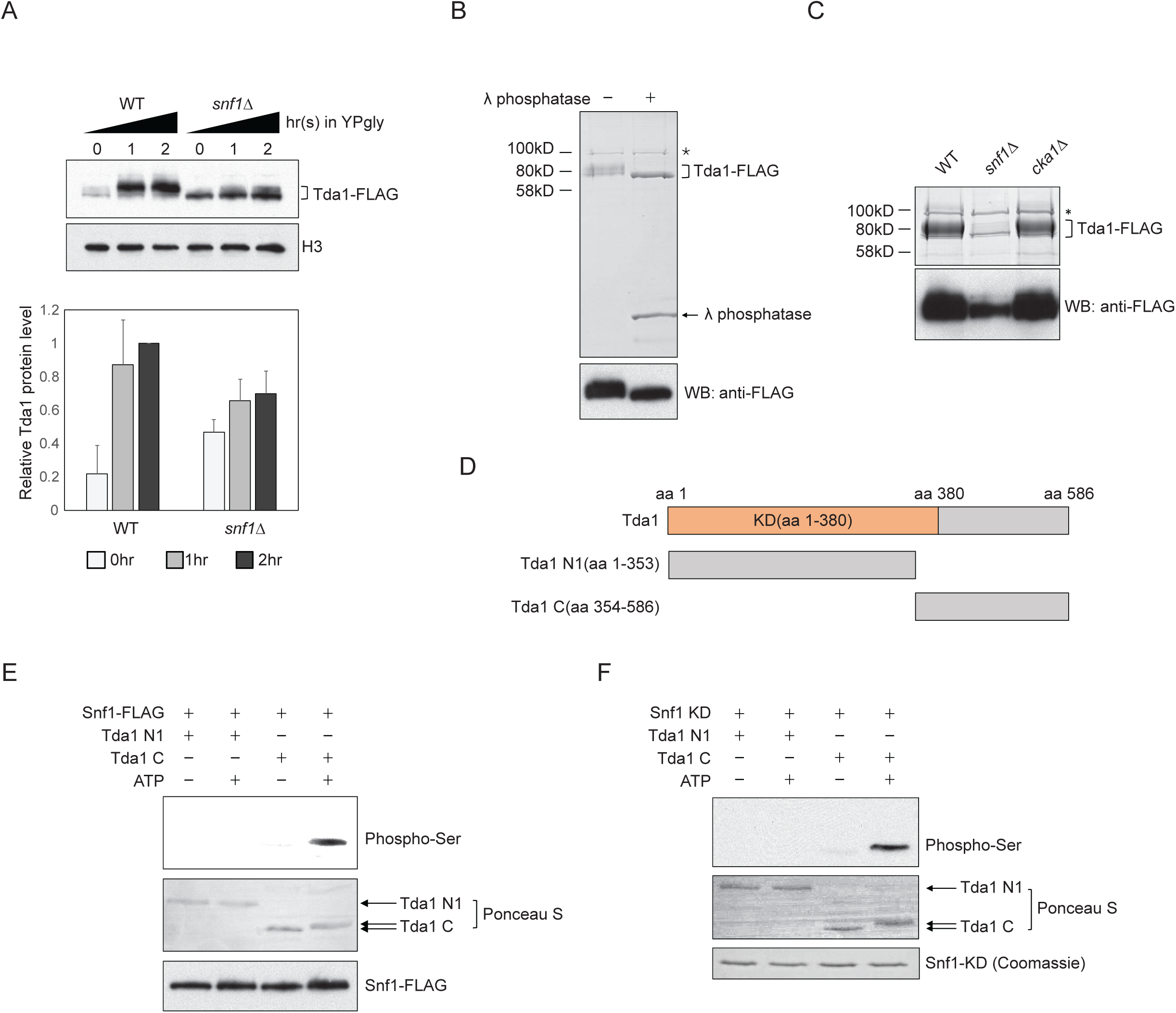
Snf1 phosphorylates Tda1 at C-terminus. (A) (Upper panel) Tda1-3xFLAG protein level changes in WT and *snf1Δ* upon media shift from YPD to YPgly measured by western blot against FLAG tag. (Lower panel) Relative band intensities of Tda1 to H3 with error bars indicating STD of three biological replicates. (B) Coomassie staining (upper panel) or western blot (WB) against FLAG tag (lower panel) of yeast FLAG purified Tda1 with or without λ phosphatase treatment. (C) Coomassie staining (upper panel) or western blot against FLAG tag (lower panel) of yeast FLAG purified Tda1 in WT, *snf1Δ*, and *cka1Δ* background. (D) Schematic diagram of Tda1 N1 (Tda1 aa 1 to 353) and Tda1 C (Tda1 aa 354 to 586) used for *in vitro* kinase assays shown in (E) and (F). (E) *In vitro* kinase assay of yeast FLAG purified Snf1 from *reg1Δ* background using GST-Tda1 N1 or GST-Tda1 C as a substrate. (F) *In vitro* kinase assay of recombinant Snf1 kinase domain (Snf1-KD) which was activated by human CaMKK2 using GST-Tda1 N1 or GST-Tda1 C as a substrate.

Tda1 co-immunoprecipitated with Snf1 as well as with Cka1 *in vivo* (Figure 5-figure supplement 1B and 1C). In this regard, we tested whether Snf1 can directly phosphorylate Tda1 *in vitro*. Snf1 was purified from a *reg1Δ* strain using FLAG tag, maintaining a robust Snf1 pT210 signal (Figure 5-figure supplement 1D), as Reg1 recruits the Snf1 pT210 phosphatase Glc7 and inactivates Snf1 (Tu & Carlson, 1995). Then the purified Snf1-FLAG was incubated with GST-tagged Tda1 fragments, Tda1 N1 (amino acids 1 to 353) or Tda1C (aa 354 to 586) (Figure 5D). Interestingly, Snf1 robustly phosphorylated only the Tda1C fragment (Figure 5E). The pattern of protein band migration in the PAGE gel also indicated phosphorylated Tda1C. An *in vitro* assay with a recombinant Snf1 kinase domain (aa 41-315, Snf1-KD) also showed that Snf1 can phosphorylate Tda1C fragment (Figure 5F).

### Tda1 phosphorylation by Snf1 is required for Tda1 activity

To investigate which residues of Tda1 are modified by Snf1, the Tda1C fragment was incubated with recombinant Snf1-KD or recombinant GST-Cka1 and then subjected to MudPIT (Multi-dimensional protein identification technology) mass spectrometry analysis. Surprisingly, MudPIT analysis revealed that Snf1 phosphorylates multiple serine and threonine residues of the Tda1C fragment (Figure 6A and Supplemental Table S1). We categorized Tda1 phosphorylation sites by Snf1 into three groups by their proximity to each other. Group I includes multiple serine and threonine residues residing in amino acids from 396 to 417. Interestingly, this region is located just after the Tda1 kinase domain. Group II includes tandem serine 483 and threonine 484. These two residues showed the highest PTM percentages among tested residues. Group III includes serine 570. MudPIT analysis also revealed that Cka1 robustly phosphorylated Tda1C at serine 578.

**Figure 6.**
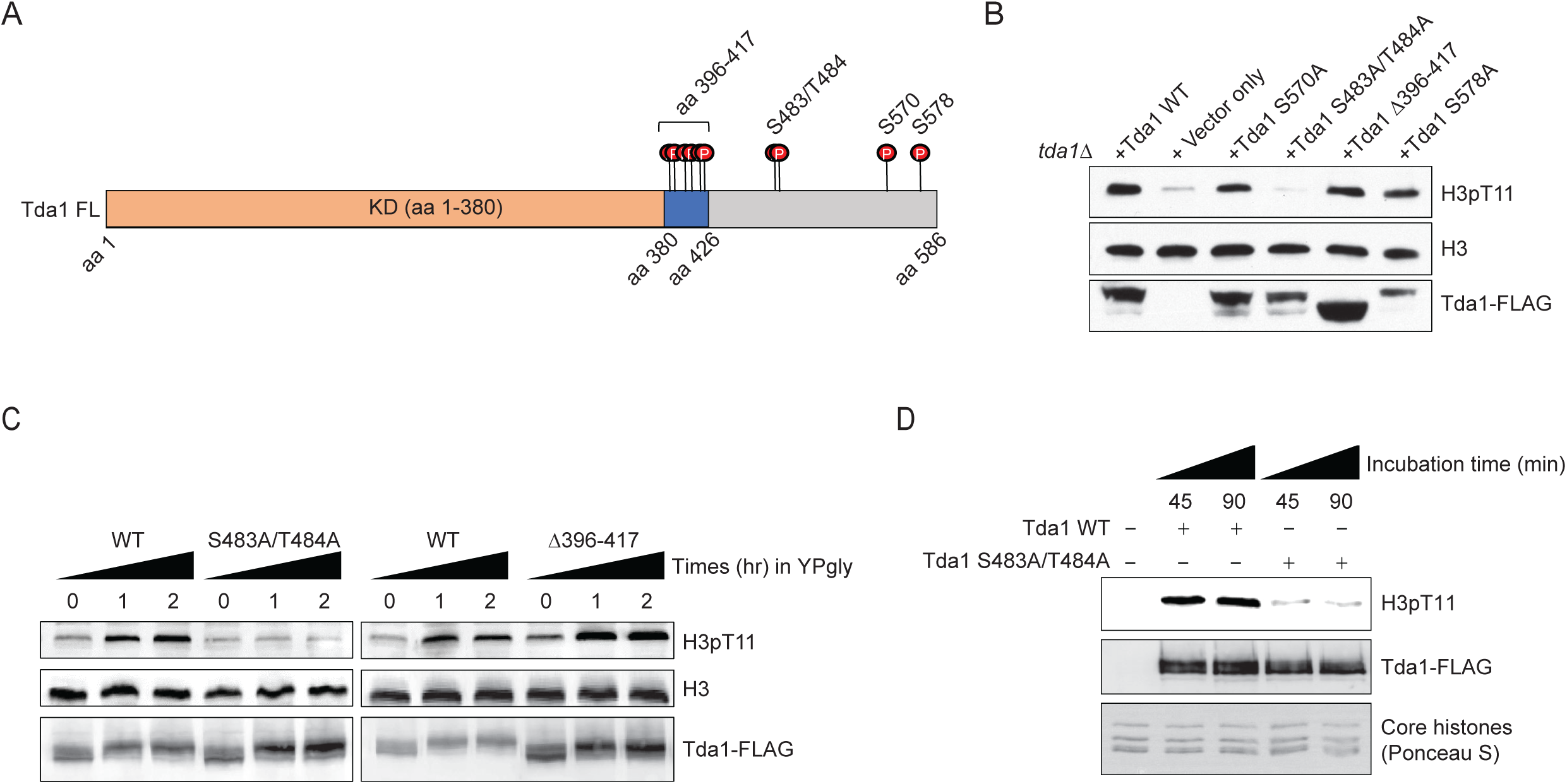
Tda1 S483/T484 phosphorylation by Snf1 is required for the Tda1 activity *in vivo*. (A) Schematic map of Tda1 phosphorylation sites (denoted as red circles) by recombinant Snf1-KD and Cka1. (B) Global H3pT11 levels of Tda1 phosphorylation sites defective mutants taken from saturated cultures in YPD media. The Tda1 constructs were expressed in pRS416 shuttling vectors and their expression was governed by ADH1 promoter. ‘Vector only’ construct indicates the pRS416 with ADH1 promoter only. (C) H3pT11 levels upon media shift from YPD to YPgly in *tda1Δ* cells expressing Tda1 WT, Tda1 S483A/T484A or Tda1 Δ396-417 constructs under an ADH1 promoter. (D) *In vitro* kinase assay of yeast FLAG purified Tda1 WT or Tda1 S483A/T484A protein using core histones as substrates.

To understand how Tda1 phosphorylation by Snf1 or Cka1 regulates its function, we generated Tda1 phosphorylation site defective mutants and tested their H3pT11 levels. Strikingly, among the tested mutants, H3pT11 level was significantly reduced only in the group II site defective mutant, Tda1 S483A/T484A (Figure 6B). Tda1 S483A/T484A mutant also showed impaired H3pT11 increase, while Tda1 Δ396-417 mutant showed intact H3pT11 increase upon media shift from YPD to YPgly (Figure 6C). Interestingly, Both Tda1 S483A/T484A and Δ396-417 mutants showed the robust increase of Tda1 phosphorylation upon media shift (Figure 6C Tda1-FLAG panel), which was significantly reduced in *snf1Δ* background (Figure 5A). These results suggest that Snf1 is responsible for the multi-sites phosphorylation of Tda1 and among those phosphorylation, S483/T484 phosphorylation has a critical role for H3pT11 regulation.

We inquired how Tda1 S483/T484 phosphorylation can regulate the function of Tda1 for H3pT11. First, we hypothesized that the availability of Tda1 for H3pT11 may be affected by its phosphorylation. However, Tda1 S483A/T484A mutant showed a similar level of nuclear Tda1 compared to Tda1 WT (Figure 6-figure supplement 1A and 1B), suggesting that Tda1 S483/T484 phosphorylation is not required for Tda1 nuclear localization. Next, we hypothesized that Tda1 S483/T484 phosphorylation may regulate Tda1 activity against H3T11. To test this hypothesis, we purified Tda1 WT or Tda1 S483A/T484A protein from yeast for *in vitro* kinase assays. Interestingly, the Tda1 S483A/T484A mutant showed significantly lower activity against H3 T11 than Tda1 WT (Figure 6D). These results indicate that Snf1 regulates Tda1 activity by phosphorylating Tda1 S483/T484 residues.

### CK2 regulates Tda1 nuclear localization

Both Snf1 and CK2 are required for proper H3pT11 levels in low glucose stress conditions, and they genetically interact with each other in H3pT11 regulation (Figure 1D). CK2 does not affect global Tda1 phosphorylation levels (Figure 5C). Tda1 purified from *cka1Δ* background showed similar activity against H3 T11 to Tda1 purified from WT (Figure 7-figure supplement 1), suggesting that CK2 does not affect Tda1 activity. To investigate how CK2 regulates Tda1 function, we tested Tda1 subcellular localization in WT and *cka1Δ* mutant.

Interestingly, we found that Tda1 nuclear localization was significantly decreased in *cka1Δ* mutants in low glucose stress conditions compared to WT (Figure 7A). As we could not find any conventional NLS sequence in Tda1, we attached a strong cMyc nuclear localization signal (PAAKRVKLD) (Dang & Lee, 1988) at the C-terminal end of Tda1 to see if the cMyc NLS could help Tda1 to bypass the requirement for Cka1 for its nuclear localization. The DNA construct expressing Tda1 tagged with C-terminal 3xFLAG (Tda1 WT) or 3xFLAG with cMyc NLS (Tda1 cNLS) under a strong ADH1 promoter was integrated into the genome of a *tda1Δ* mutant. Upon low glucose stress, Tda1 cNLS showed a slightly lower level of the protein increase than Tda1 WT. However, the Tda1 cNLS expressing strain showed a more robust and rapid increase of H3pT11 compared to the Tda1 WT expressing strain (Figure 7B). Next, we integrated the Tda1 WT or Tda1 cNLS construct into *tda1Δcka1Δ* or *tda1Δsnf1Δ* backgrounds. Interestingly, in the *cka1Δ* mutant, the Tda1 cNLS was able to mediate H3pT11, unlike Tda1 WT (Figure 7C, left panel). On the contrary, neither Tda1 cNLS nor Tda1 WT could restore H3pT11 in *snf1Δ* mutant (Figure 7C, right panel). Thus, while Snf1 regulates Tda1 activity Cka1 regulates its nuclear localization. These results clearly show that Snf1 and Cka1 signaling pathways cooperatively regulate H3pT11 but use different mechanisms or act in different stages of the Tda1 regulation (Summarized in Figure 7D).

**Figure 7.**
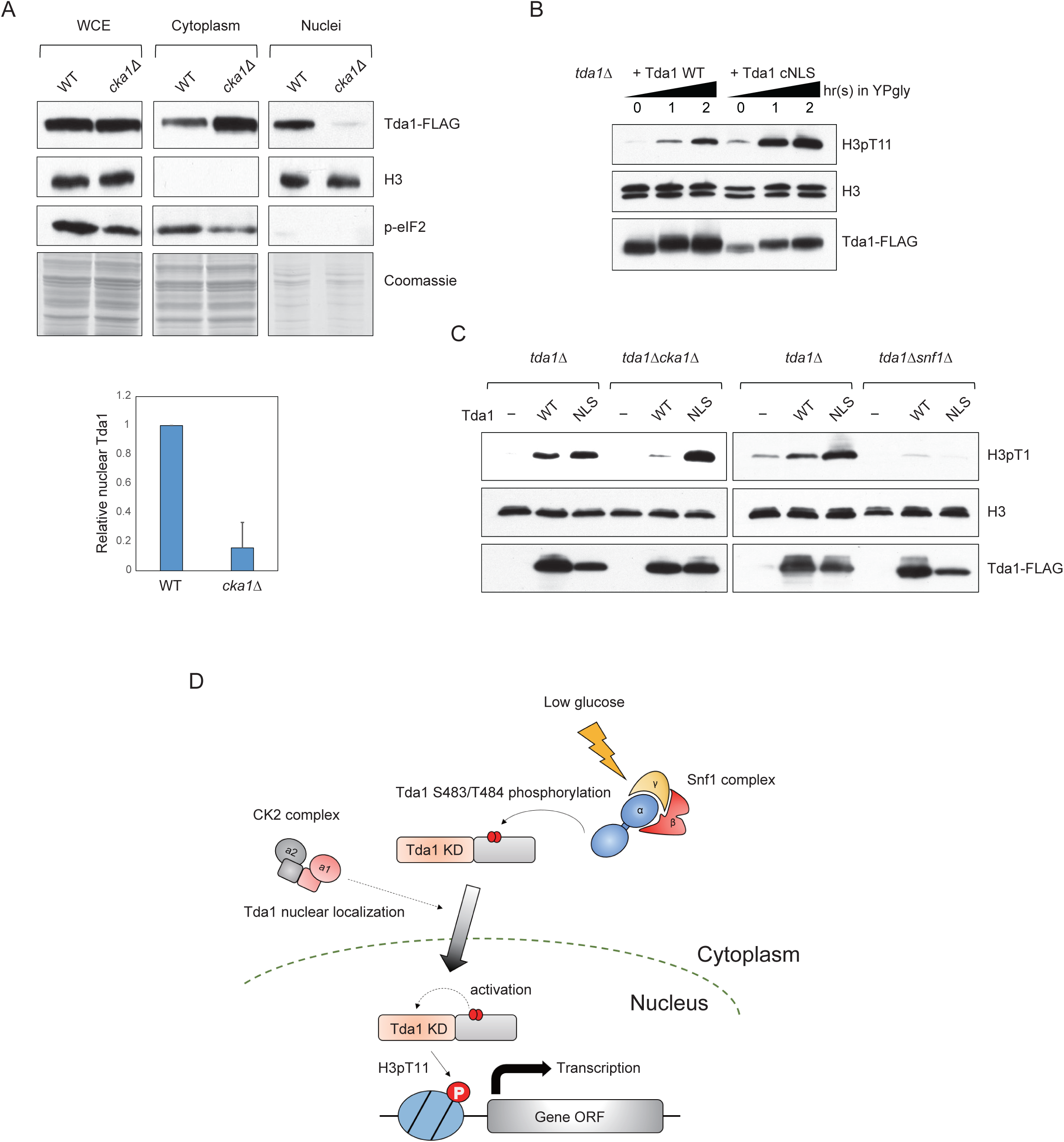
CK2 regulates Tda1 nuclear localization. (A) (Upper panels) Subcellular localization of Tda1-3xFLAG proteins in WT and *cka1Δ* mutant. WCE, cytoplasm, and nuclei samples were taken from YPgly media cultures incubated for 2 hours after the media shift from YPD, then analyzed by western blot. Histone H3 and p-eIF2 antibodies were used for the nuclear and cytoplasmic marker, respectively. (Lower panel) The relative band intensities of nuclear Tda1 to WCE Tda1 are presented with error bars indicating STD of three biological replicates. (B) H3pT11 level changes in Tda1 WT and Tda1 with C-terminally tagged cMyc NLS upon media shift from YPD to YPgly analyzed by western blots. (C) H3pT11 restoration by genome integrated Tda1 WT or Tda1 with cMyc NLS in *tda1Δ, tda1Δcka1Δ*, and *tda1Δsnf1Δ* backgrounds measured by western blots. The cells were taken from YPgly media cultures incubated for 2 hours after the media shift from YPD. (D) Summary model of the Tda1 regulation by Snf1 and Cka1. The red ellipses represent phosphorylation.

## Discussion

### Tda1 phosphorylation by Snf1

*In vitro* kinase assays using recombinant Tda1 fragments and Snf1 revealed multiple Tda1 residues phosphorylated by Snf1 (Figure 6A and Supplemental Table S1). Among Tda1 phosphorylation site-defective mutants tested, only a Tda1 S483A/T484A mutant showed a significant defect in H3pT11 regulation *in vivo* (Figures 6B and C), suggesting the importance of Tda1 S483/T484 phosphorylation. Indeed, the Tda1 S483/T484 residues showed the highest level of modification in the *in vitro* kinase assay (Supplemental Table S1).

Tda1 S483/T484 phosphorylation is required for Tda1 kinase activity on H3 T11 (Figure 6D), however, How the modification regulates Tda1 activity remains unclear. Tda1 S483/T484 residues are not located in the Tda1 catalytic domain (Figure 4E, Tda1 aa 1-380). Tda1 S483/T484 phosphorylation may regulate Tda1 activity by relieving an unknown inhibition mechanism, or Tda1 may require additional factors to become fully active, and Tda1 phosphorylation at S483/T484 may be responsible for interacting with such factors. Investigating Tda1 binding proteins which depend on Tda1 phosphorylation would be an exciting future step.

### Cka1 regulates Tda1 nuclear localization

Both Snf1 and Cka1 are required for H3pT11 regulation, but two kinases regulate H3pT11 via different mechanisms. Snf1 robustly phosphorylates the Tda1 C terminal (Figures 5E and F), while Cka1 does not affect global Tda1 phosphorylation (Figure 5C). A Tda1 S578A point mutation at the putative phosphorylation site by Cka1 does not affect the global level of H3pT11 (Figure 6B). Tda1 with cMyc NLS behaves differently in the *snf1Δ* and *cka1Δ* backgrounds (Figure 7C). The NLS-attached Tda1 can bypass the requirement for Cka1, but not that for Snf1. This result suggests that Cka1 regulates the nuclear localization process of Tda1 by controlling the interaction between Tda1 and nuclear importins, in contrast to Snf1 which regulates Tda1 enzymatic activity (Figure 6D). Tda1 does not possess conventional phosphorylation NLS sequence in it. In this regard, it is not clear if Tda1 itself contains non-conventional NLS, or Tda1 requires other accessory factors for its nuclear localization. Finding a Tda1 specific importin would also be an important next step.

CK2 has been known as a constitutively active kinase complex, which does not require upstream stimulation for its activity. However, many processes governed by CK2 are highly controlled processes (Pinna, 2002). In this study, we show an example of how a constitutively active kinase can be involved in tightly regulated processes such as H3pT11 upon low glucose stress. Although CK2 is constitutively active, H3pT11 cannot be achieved without active Snf1. Previously, we reported that Sch9 was also required for H3pT11 regulation as well as CK2 (Oh et al., 2018). Interestingly, Sch9 and CK2 genetically interact in H3pT11 regulation as Cka1 and Snf1 do. Our finding of H3pT11 regulation by CK2 via controlling the Tda1 nuclear localization suggests a possibility that Sch9 is also involved in the same process. This raises the possibility of Tda1 being a signaling platform that combines multiple signaling such as Snf1, CK2, and Sch9 together to finely connect external glucose levels to chromatin.

### Tda1 in higher eukaryotes

Structural analysis (at http://yeastgenome.org) predicts that Tda1 contains a calcium/calmodulin dependent kinase domain at its N-terminus. We found that its functional kinase domain resides between amino acid 1 to 380 (Figure 4F). In yeast, there are 12 kinases that contain the domain including yeast calcium calmodulin dependent kinases (Cmk1 and Cmk2) and meiosis specific kinase Mek1, as well as Tda1. Interestingly, Mek1 phosphorylates H3 at T11 during meiosis (Kniewel et al., 2017). A large scale screening of yeast phosphorylation site motifs revealed that the amino acid sequence surrounding H3T11 fits in the recognition motif of Cmk1 and Cmk2 (Mok et al., 2010). These results imply that calcium/calmodulin dependent kinases could be kinase candidates for H3pT11 in situations other than nutrient starvation in yeast and possibly in other species.

Based on predictions by the PPOD program (at http://ppod.princeton.edu/), Nuak1 has been proposed as a human ortholog of yeast Tda1 (Soma, Yang, Morales, & Polymenis, 2014). Nuak1 is also known as ARK5 which means AMPK related kinase 5. As its name suggests, Nuak1 has a 47% amino acid sequence homology to AMPK α1 (Suzuki, Kusakai, Kishimoto, Lu, Ogura, Lavin, et al., 2003). Nuak1 is involved in cell survival processes upon nutrient starvation (Suzuki, Kusakai, Kishimoto, Lu, Ogura, & Esumi, 2003), suggesting the Tda1 function upon nutrient stress shown in yeast is consistent in higher eukaryotes. Nuak1 predominantly localizes to the nucleus, but a recent study revealed that the cellular distribution of Nuak1 changes upon stresses in a importin β-dependent manner (Palma et al., 2019). Investigating whether the Nuak1 subcellular localization is regulated via a similar mechanism to that of Tda1 and phosphorylates histones under a specific stress like Tda1 would be an interesting future study.

## Materials and Methods

### Yeast strains and culture conditions

All yeast strains used in this study are listed in Table S2. All single deletion mutants using KanMX4 cassette and TAP tagged strains derived from BY4741 were obtained from Open Biosystems library (maintained at the Stowers Institute Molecular Biology facility). Yeast synthetic histone H3 mutants (Dai et al., 2008) (H3 WT, H3 T11A and H3 S10A in Figure 1F) were also purchased from Open Biosystems. Other deletions and tagged strains were made by targeted homologous recombination of PCR fragments containing marker genes flanked by gene specific sequences. These strains were confirmed by PCR with primer sets specific for their marker genes. For Tda1 mutant strains shown in Figures 6C, 6D and 7, Tda1 WT or Tda1 mutant constructs were cloned into pRS406 vector and the DNA constructs were linearized by NcoI digestion then integrated into URA3 loci of the genome. For low glucose stress experiments, yeast cells were saturated by overnight culture at 30°C. For media shift experiments, the saturated cultures were inoculated into fresh YPD media and then incubated until mid-log phase (Optical density 0.4-0.6). These cultures were washed once with YPgly (YP with 3% glycerol) media, then resuspended with YPgly for indicated times at 30°C.

### Preparation of yeast whole cell extracts

Yeast whole cell extracts were prepared as previously described (Oh et al., 2018). 5 OD cells were taken from the cultures. Cell pellets were washed once with distilled water, then resuspended in 250 uL of 2M NaOH with 8 % β-Mercaptoethanol for 5 minutes on ice. Cells were pelleted then washed once with 250 uL TAP extraction buffer (40 mM HEPES pH 7.5, 10% Glycerol, 350 mM NaCl, 0.1% Tween-20, phosphatase inhibitor cocktail and proteinase inhibitor cocktail from Roche). Cell pellets were resuspended in 180 uL modified 2X SDS buffer then boiled at 100 °C for 4-5 minutes. For detecting Snf1 T210 phosphorylation (Figure 1A), cell cultures were boiled for 5 minutes before initial harvesting to prevent Snf1 phosphorylation by centrifugation (Orlova, Barrett, & Kuchin, 2008).

### Yeast flag tagged protein purification

Tda1 and Snf1 proteins with 3xFLAG tag were purified as previously described (Dutta et al., 2014) with minor modifications. 6L cultures of cells were grown in YPD to 2-3 OD at 30°C and collected. For Tda1-3xFLAG purification, cell pellets were resuspended in 3L YP without glucose and then incubated for an additional 1 hour. Cell pellets were washed once with distilled water, then resuspended in buffer A (25 mM HEPES pH 7.5, 10% glycerol, 350 mM KCl, 2 mM MgCl_2_, 1 mM EDTA, 0.02% NP40, supplemented with 20 µg/mL leupeptin, 20 µg/mL pepstatin and 100 µM PMSF) and then broken up by bead beating. The crude cell extracts were incubated with 125U Benzonase and 500 µg heparin for 15 minutes at RT to remove nucleic acid contamination, then the extracts were further clarified by ultracentrifugation. The clarified extracts were incubated with Anti-FLAG M2 affinity resin (Sigma) for 4 hours at 4 °C with gentle rotation. Proteins bound to the resin were washed 3 times with Buffer A, then washed once with Buffer B (25 mM HEPES pH 7.5, 10% glycerol, 100 mM KCl, 2 mM MgCl_2_, 1 mM EDTA, 1 mM DTT, 0.02% NP40, supplemented with 20 µg/mL leupeptin, 20 µg/mL pepstatin and 100 µM PMSF). Bound proteins were eluted in buffer B containing 0.5 mg/mL 3xFLAG peptides.

### Recombinant protein purification

Yeast Cka1 and Tda1 genes were amplified by PCR from yeast genomic DNA and then cloned into pGEX4T-1 (GE healthcare) vector. GST-Snf1 kinase domain (Snf1-KD) expression vector was purchased from Addgene (#52683). Those DNA constructs were transformed in Rosetta2 (Novagen) competent cells and protein expressions were induced by 0.5 mM IPTG for 18 hours at 16°C. Bacterial cell pellets were resuspended in TAP extraction buffer, then sonicated to disrupt cell walls. Crude extracts were incubated with Glutathione sepharose 4B (GE healthcare) resin for 3 hours at 4°C. Resin bound proteins were eluted in 50 mM Tris pH 8.0 buffer containing 20 mM Glutathione. *Xenopus* core histones were purified as previously described (Shim, Duan, Chen, Smerdon, & Min, 2012) with minor modifications. Briefly, YS-14 construct encoding all four *Xenopus* core histones (H2A, H2B, H3, and H4) was transformed into Tuner DE3 pLysS competent cells (Novagen), and then histone protein expressions were induced in 2X YT media by 0.5 mM IPTG for 24 hours at 37°C. Cell pellets were sonicated in high salt extraction buffer (20 mM Tris pH 8.0, 2M NaCl), and then clarified extracts were incubated with TALON metal affinity resin (Clontech) for 3 hours at 4°C. Resin bound proteins were eluted by 250 mM imidazole, then eluted proteins were dialyzed in high salt extraction buffer for overnight at 4°C.

### Yeast subcellular fractionation

Yeast cellular fractionation was carried out as previously described (Emili et al., 2006) with minor modifications. 40 to 50 OD yeast cells were pelleted, then washed successively with 10 mL distilled water and ice cold 10 mL SB (1 M Sorbitol, 20 mM Tris-Cl pH 7.5), The washed cell pellets were transferred to 1.7 mL Eppendorf tube, then successively washed with 1 mL PSB (20 mM Tris-Cl pH 7.5, 2 mM EDTA, 100 mM NaCl, 10 mM β-Mercaptoethanol) and 1mL SB. The pellets were resuspended with 750 µL SB, then yeast cell walls were digested by adding 100 µL Zymolyase (10 mg/mL, Seikagaku) for 1 hour at RT. After the cell wall digestion, 750 µL ice-cold SB was added, then the spheroplasts were collected by gentle centrifugation (2K, 5 mins, 4°C) and washed once with 1 mL ice-cold SB. The spheroplasts were resuspended by 500 µL EBX (20 mM Tris-Cl pH 7.5, 100 mM NaCl, 0.25 % Triton X-100, 15 mM β-Mercaptoethanol), then 100 % Triton X-100 was added up to 0.5% to disrupt outer cell membrane. Cells were placed on ice for 10 minutes with occasional mixing, then 1 mL NIB (20 mM Tris-Cl pH 8.0, 100 mM NaCl, 1.2 M Sucrose, 15 mM β-Mercaptoethanol) was layered over the cells. After high speed centrifugation (12K, 15 mins, 4°C), the upper layer was taken as cytoplasmic fraction. Glassy, white nuclear pellets were resuspended SDS containing sample buffer then boiled for western blot experiments.

### In vitro kinase assays

All *in vitro* kinase assays were done in NEBuffer for protein kinase (NEB) at 30°C. Recombinant Snf1 kinase domain (Snf1-KD) was phosphorylated by recombinant human CaMKK2 (Abnova) for its activation before kinase assays (Figure 5F). When Tda1 fragments were used as kinases (Figure. 4F), additional DTT was supplemented up to 1 mM. The reactions were quenched by SDS sample buffer addition, then boiled at 100°C for 5 minutes.

### Multi-dimensional Protein Identification Technology (MudPIT) analysis

TCA-precipitated proteins were urea-denatured, reduced, alkylated and digested with endoproteinase Lys-C (Promega) followed by modified trypsin (Promega) as previously described (Florens & Washburn, 2006; Washburn, Wolters, & Yates, 2001). Peptide mixtures were loaded onto 100 µm fused silica microcapillary columns packed with 5-μm C_18_ reverse phase (Aqua, Phenomenex), strong cation exchange particles (Partisphere SCX, Whatman), and reverse phase (MacCoss et al., 2002). Loaded microcapillary columns were placed in-line with a Quaternary Agilent 1260 series HPLC pump and a Velos Orbitrap mass spectrometer equipped with a nano-LC electrospray ionization source (ThermoFinnigan). Fully automated 10-step MudPIT runs were carried out on the electrosprayed peptides, as previously described (Florens & Washburn, 2006). Tandem mass (MS/MS) spectra were interpreted using ProLuCID (Eng, McCormack, & Yates, 1994) against a database of 8956 sequences, consisting of 4303 *E. coli* proteins (downloaded from NCBI on 2013-11-06), 177 usual contaminants such as human keratins, IgGs, and proteolytic enzymes, 4476 “shuffled” sequences and four yeast proteins sequences (Tda1, Snf1, Cka1 and a recombinant Tda1 C-terminal sequence from amino acid residues 354 to 586). Peptide/spectrum matches were sorted and selected using DTASelect (Tabb, McDonald, & Yates, 2002) with the following criteria set: spectra/peptide matches were only retained if they had a DeltCn of at least 0.08, and minimum XCorr of 1.8 for singly-, 2.0 for doubly-, and 3.0 for triply-charged spectra. In addition, peptides had to be fully tryptic and at least 6 amino acids long. Combining all runs, proteins had to be detected by at least 2 such peptides, or 1 peptide with 2 independent spectra. Under these criteria, the final FDRs at all levels were zero. Peptide hits from multiple runs were compared using CONTRAST (Tabb et al., 2002). To estimate relative protein levels, distributed Normalized Spectral Abundance Factors (dNSAFs) were calculated for each detected protein, as previously described (Florens et al., 2006; Paoletti et al., 2006; Zybailov et al., 2006).

## Acknowledgements

We thank Dr. Michael Church for scientific editing of the manuscript. We thank the Workman Lab members and Stowers core facilities for support during this project. These studies were supported by funds from the Stowers Institute and the National Institutes of General Medical Sciences grant R35GM118068 (J.L.W.). Original data underlying this manuscript can be accessed from the Stowers Original Data Repository at http://www.stowers.org/research/publications/libpb-1536.

## Author contributions

S.O. and J.L.W. designed the study, analyzed the data, and wrote the manuscript. S.O. and J.L. performed the experiments. S.K.S., L.P, and M.P.W. analyzed the proteomics data.

## Figure legends

**Figure 1-figure supplement 1.**
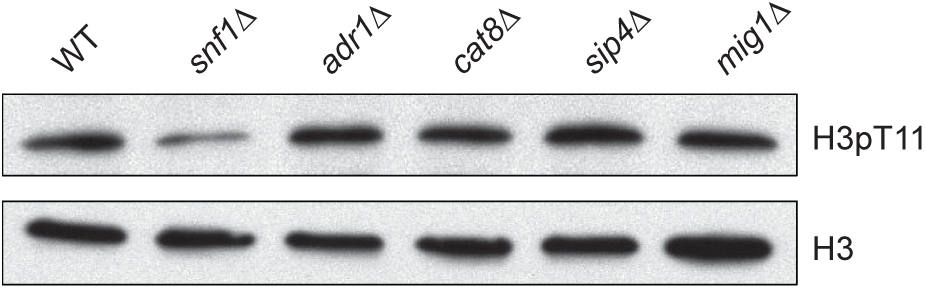
H3pT11 depends on Snf1, but not on Snf1 target transcription factors. Global H3pT11 levels in WT (BY4741), *snf1Δ*, and the deletion mutants of Snf1 target transcription factors (*adr1Δ, cat8Δ, sip4Δ* and *mig1Δ*) analyzed by western blots. The cells were taken from overnight saturated cultures in YPD media.

**Figure 3-figure supplement 1.**
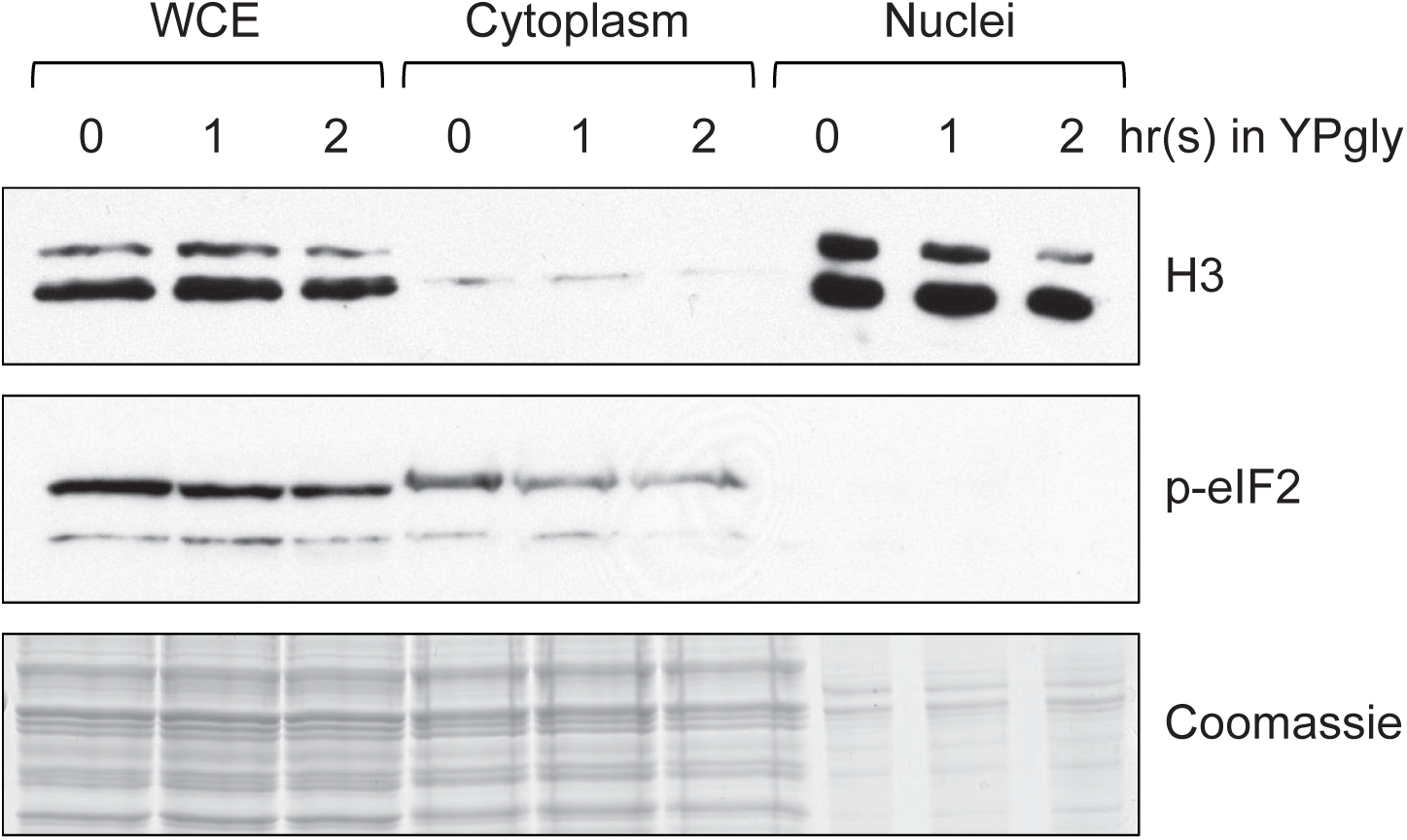
Confirmation of efficient yeast subcellular fractionation of the samples shown in Figure 3D. 1% of whole cell extract (WCE), 1% of cytoplasm, and 2% of nuclei samples used in Figure 3D were subjected to western blots to confirm efficient subcellular fractionation. Histone H3 and phosphorylated eIF2 (p-eIF2) antibodies were used for the nuclear and cytoplasmic marker, respectively.

**Figure 5-figure supplement 1.**
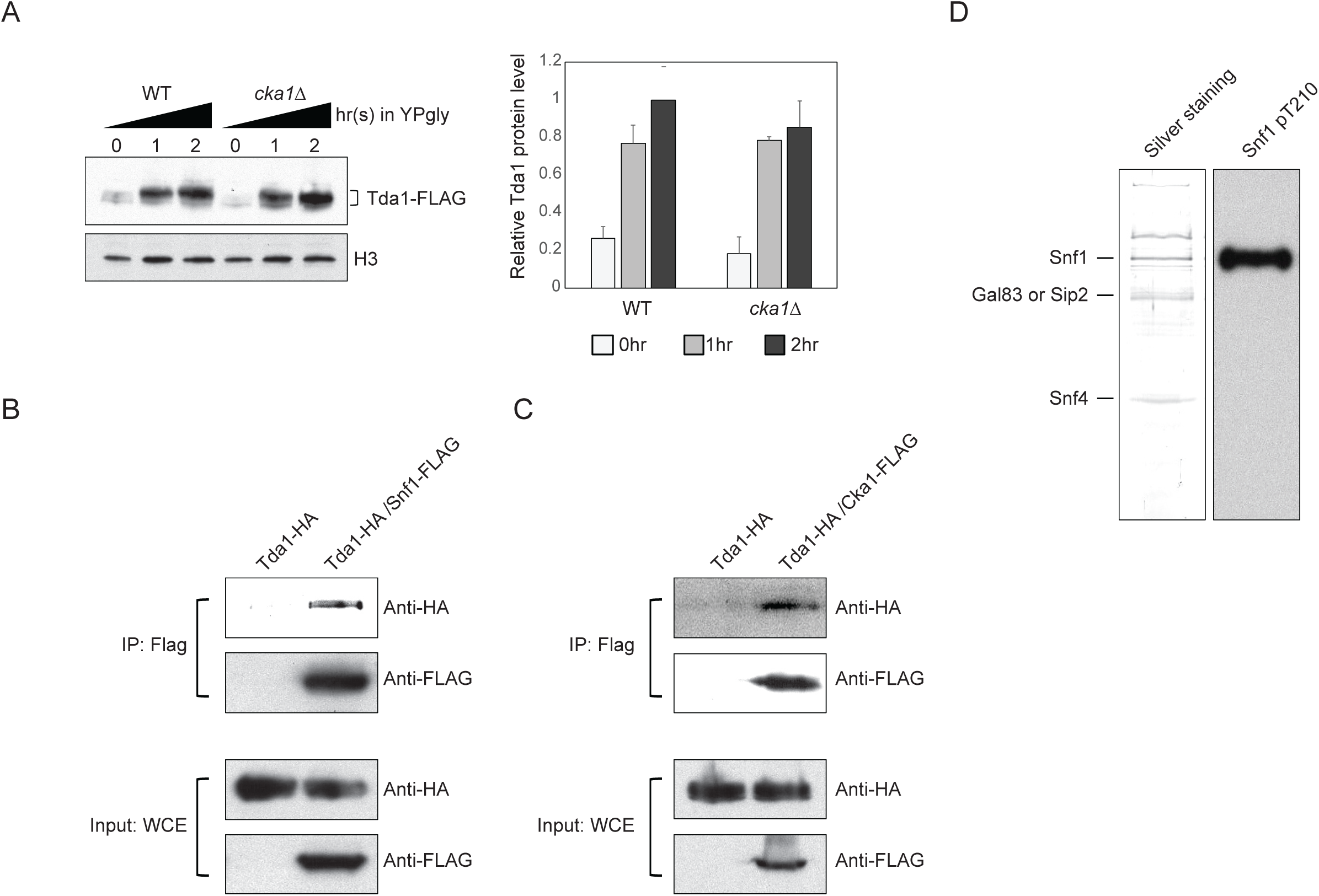
Snf1 phosphorylates Tda1 at C terminus. (A) (Left panel) Tda1-3xFLAG protein level changes in WT and *cka1Δ* upon media shift from YPD to YPgly measured by western blot against FLAG tag. (Right panel) The relative band intensities of Tda1 to H3 with error bars indicating STD from three biological replicates. (B and C) Tda1 co-immunoprecipitation assay using (B) FLAG tagged Snf1 or (C) FLAG tagged Cka1 as a bait in YPgly media. Immunoprecipitated Tda1 was detected by western blot against HA tag. Tda1-HA strain without any FLAG tagged protein was used as a negative control. (D) The silver staining (left panel) or western blot against Snf1 pT210 (right panel) of yeast FLAG purified Snf1 from *reg1Δ* background.

**Figure 6-figure supplement 1.**
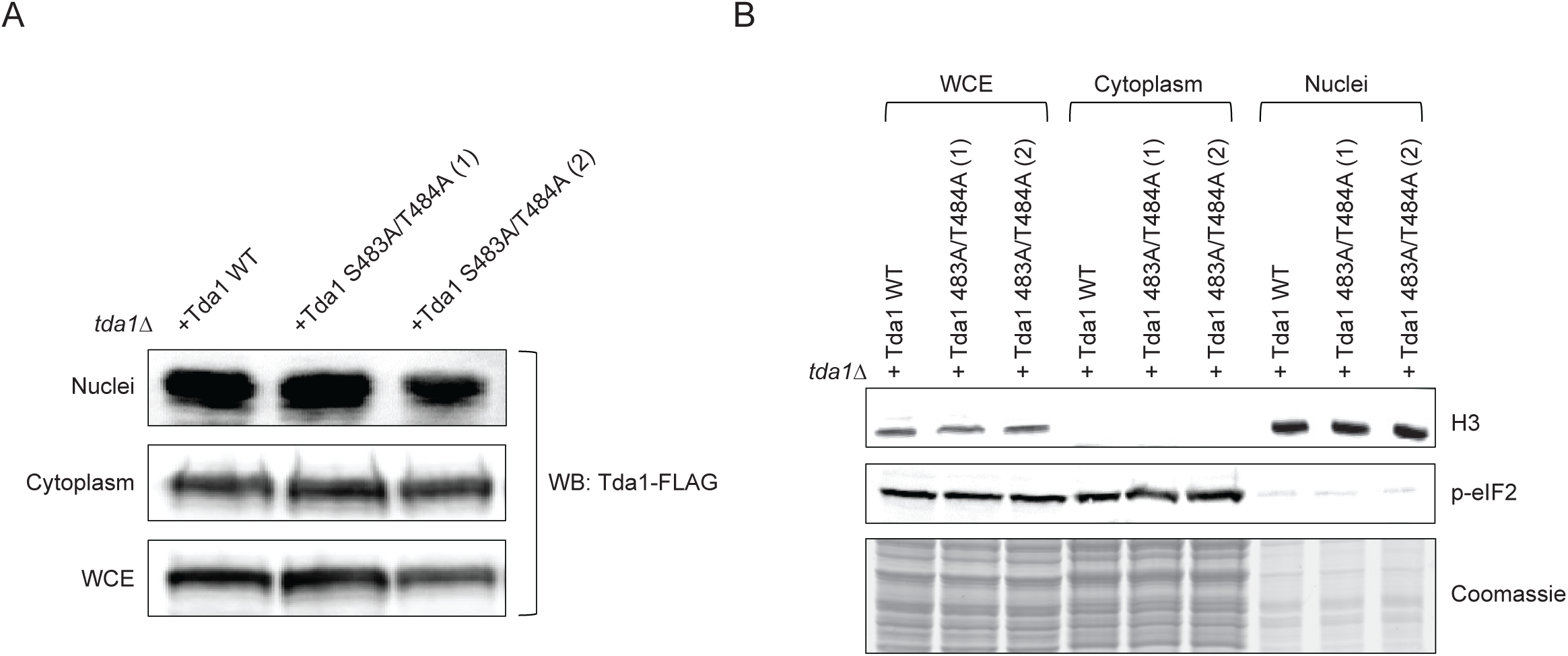
Tda1 S483/T484 phosphorylation by Snf1 is required for the nuclear Tda1 activity. (A) Subcellular localization of Tda1 WT and Tda1 S483A/T484A proteins tagged with C terminal 3xFLAG tag measured by western blot. The cells were taken from YPgly media cultures incubated for 2 hours after the media shift from YPD. (B) 0.5% of whole cell extract (WCE), 0.5% of cytoplasm, and 1% of nuclei samples used in (A) were subjected to western blots to confirm efficient subcellular fractionation. Histone H3 and p-eIF2 antibodies were used for the nuclear and cytoplasmic marker, respectively.

**Figure 7-figure supplement 1.**
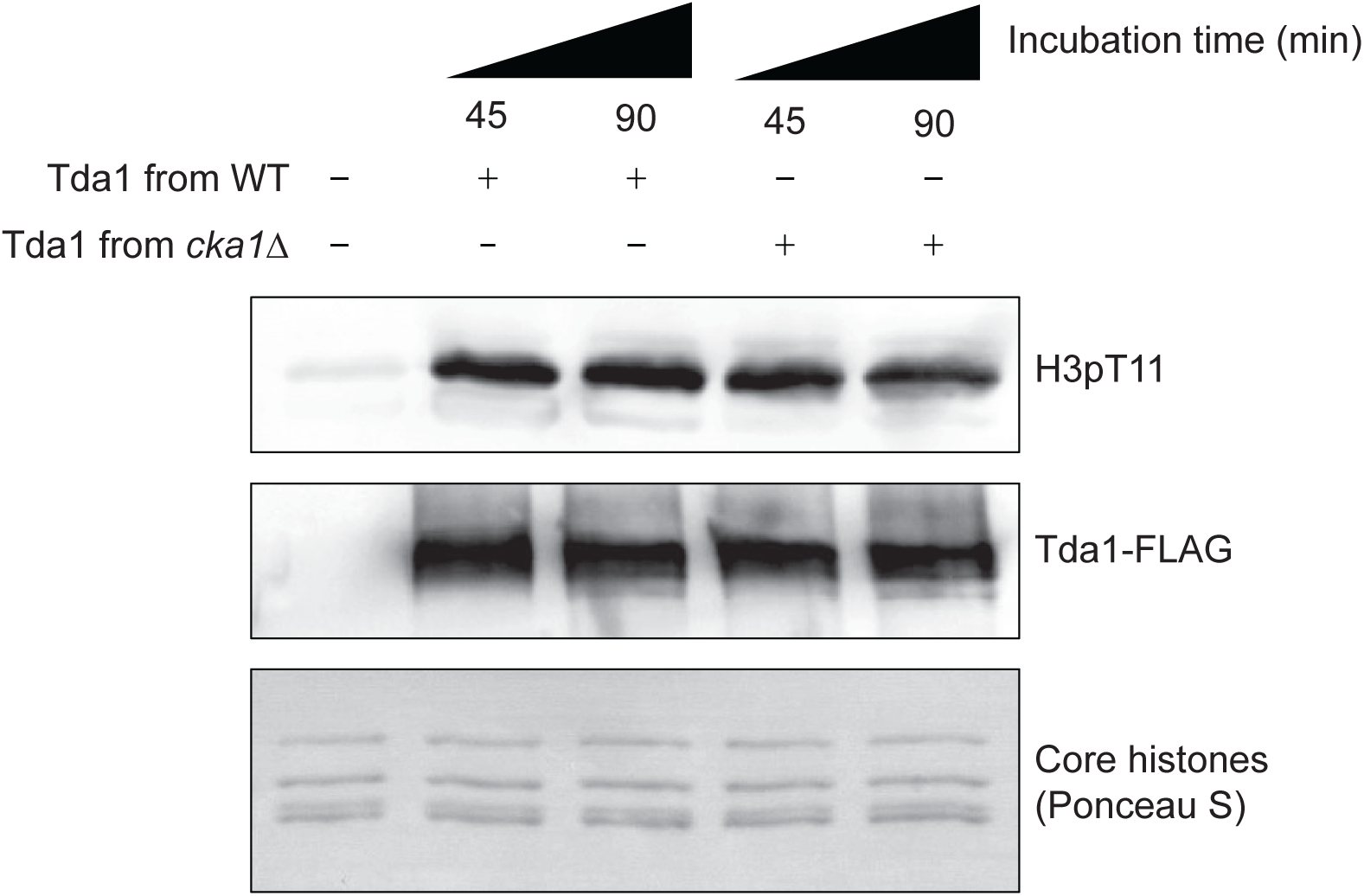
CK2 does not regulate Tda1 activity. *In vitro* kinase assay of yeast FLAG tagged Tda1 purified from WT or *cka1Δ* background. Core histones were used as substrates.

