## Supplemental tables 1 and 2 for "Yeast Nuak1 phosphorylates histone H3 threonine 11 in low glucose stress conditions by the cooperation of AMPK and CK2 signaling"

| Group | | Tda1C only | | | Tda1C + rSnf1-KD | | | Tda1C + rCka1 | | |
|---|---|---|---|---|---|---|---|---|---|---|
| | Residue | Total | Modified | PTM % | Total | Modified | PTM % | Total | Modified | PTM % |
| Group I | S393 | 16 | 0 | 0 | 418 | 31 | 7.42 | 7 | 0 | 0 |
| Group I | T396 | 4550 | 0 | 0 | 1449 | 203 | 14.0097 | 4808 | 0 | 0 |
| Group I | T397 | 4550 | 0 | 0 | 1449 | 69 | 4.76191 | 4808 | 0 | 0 |
| Group I | T398 | 4550 | 0 | 0 | 1449 | 151 | 10.421 | 4808 | 0 | 0 |
| Group I | Y401 | 4550 | 0 | 0 | 1449 | 65 | 4.48585 | 4808 | 0 | 0 |
| Group I | S408 | 4550 | 0 | 0 | 1449 | 68 | 4.69289 | 4808 | 0 | 0 |
| Group I | S409 | 4550 | 0 | 0 | 1449 | 56 | 3.86473 | 4808 | 0 | 0 |
| Group I | T411 | 4550 | 0 | 0 | 1449 | 202 | 13.9407 | 4808 | 0 | 0 |
| Group I | S412 | 4550 | 0 | 0 | 1449 | 105 | 7.24638 | 4808 | 0 | 0 |
| Group I | T413 | 4550 | 0 | 0 | 1449 | 164 | 11.3182 | 4808 | 0 | 0 |
| Group I | S416 | 4550 | 0 | 0 | 1449 | 337 | 23.2574 | 4808 | 0 | 0 |
| Group I | S417 | 4550 | 0 | 0 | 1449 | 297 | 20.4969 | 4808 | 0 | 0 |
| Group II | S483 | 433 | 0 | 0 | 520 | 388 | 74.6154 | 780 | 0 | 0 |
| Group II | T484 | 433 | 0 | 0 | 520 | 388 | 74.6154 | 780 | 0 | 0 |
| Group III | S570 | 2414 | 0 | 0 | 1477 | 500 | 33.8524 | 5140 | 0 | 0 |
| | S578 | 2414 | 0 | 0 | 1477 | 0 | 0 | 5140 | 4991 | 97.1 |

**Supplemental Table S1. Tda1 C fragment phosphorylation by recombinant Snf1 KD and Cka1.** The MudPIT analysis results showing the Tda1 C fragment (Tda1C, Tda1 aa 354 to 586) phosphorylation sites from *in vitro* kinase assays using Tda1C only, Tda1C with Snf1-KD (Tda1C + rSnf1-KD), and Tda1C with Cka1 (Tda1C + rCka1), respectively. Snf1-KD was activated by pre-incubation with human CaMKK2 before the kinase assay with Tda1C. The Tda1 phosphorylation sites by Snf1-KD are classified into 3 groups (I, II, and III) by the proximity of phosphorylation sites. (Total: Total peptides detected, Modified: The number of phosphorylation containing peptides, PTM%: The percentage of modified peptide compared to total peptide detected.)

| Name | Genotype | Source |
|---|---|---|
| BY4741 | *MATa his3Δ1 leu2Δ0 met15Δ0 ura3Δ0* | Open Biosystems |
| ySE18 | *MATa his3Δ1 leu2Δ0 met15Δ0 ura3Δ0 cka1Δ::KanMX4* | Open Biosystems |
| y1171 | *MATa his3Δ1 leu2Δ0 met15Δ0 ura3Δ0 tda1Δ::KanMX4* | Open Biosystems |
| y1084 | *MATa his3Δ1 leu2Δ0 met15Δ0 ura3Δ0 rim15Δ::KanMX4* | Open Biosystems |
| y1085 | *MATa his3Δ1 leu2Δ0 met15Δ0 ura3Δ0 snf1Δ::KanMX4* | Open Biosystems |
| y1129 | *MATa his3Δ1 leu2Δ0 met15Δ0 ura3Δ0 gal83Δ::KanMX4* | Open Biosystems |
| y1130 | *MATa his3Δ1 leu2Δ0 met15Δ0 ura3Δ0 sip1Δ::KanMX4* | Open Biosystems |
| y1131 | *MATa his3Δ1 leu2Δ0 met15Δ0 ura3Δ0 sip2Δ::KanMX4* | Open Biosystems |
| y1132 | *MATa his3Δ1 leu2Δ0 met15Δ0 ura3Δ0 snf4Δ::KanMX4* | Open Biosystems |
| *adr1Δ* | *MATa his3Δ1 leu2Δ0 met15Δ0 ura3Δ0 adr1Δ::KanMX4* | Open Biosystems |
| *cat8Δ* | *MATa his3Δ1 leu2Δ0 met15Δ0 ura3Δ0 cat8Δ::KanMX4* | Open Biosystems |
| *sip4Δ* | *MATa his3Δ1 leu2Δ0 met15Δ0 ura3Δ0 sip41Δ::KanMX4* | Open Biosystems |
| *mig1Δ* | *MATa his3Δ1 leu2Δ0 met15Δ0 ura3Δ0 mig1Δ::KanMX4* | Open Biosystems |
| y1163 | *MATa his3Δ1 leu2Δ0 met15Δ0 ura3Δ0 snf1Δ::KanMX4 cka1Δ::NatMX6* | In this study |
| y1166 | *MATa his3Δ200 leu2Δ0 lys2Δ0 trp1Δ63 ura3Δ0 met15Δ0 can1::MFA1pr-HIS3 hht1-hhf1::NatMX4 hht2-hhf2::[HHTS-HHFS]*-URA3* | Open Biosystems |
| y1167 | *MATa his3Δ200 leu2Δ0 lys2Δ0 trp1Δ63 ura3Δ0 met15Δ0 can1::MFA1pr-HIS3 hht1-hhf1::NatMX4 hht2-hhf2::[HHTS T11A-HHFS]*-URA3* | Open Biosystems |
| y1169 | *MATa his3Δ200 leu2Δ0 lys2Δ0 trp1Δ63 ura3Δ0 met15Δ0 can1::MFA1pr-HIS3 hht1-hhf1::NatMX4 hht2-hhf2::[HHTS S10A-HHFS]*-URA3* | Open Biosystems |
| y2242 | *MATa his3Δ1 leu2Δ0 met15Δ0 ura3Δ0 cka1-TAP::HIS3* | Open Biosystems |
| *ctk1Δ* | *MATa his3Δ1 leu2Δ0 met15Δ0 ura3Δ0 ctk1Δ::KanMX4* | Open Biosystems |
| *dbf2Δ* | *MATa his3Δ1 leu2Δ0 met15Δ0 ura3Δ0 dbf2Δ::KanMX4* | Open Biosystems |
| *elm1Δ* | *MATa his3Δ1 leu2Δ0 met15Δ0 ura3Δ0 elm1Δ::KanMX4* | Open Biosystems |
| *fus3Δ* | *MATa his3Δ1 leu2Δ0 met15Δ0 ura3Δ0 fus3Δ::KanMX4* | Open Biosystems |
| *gcn2Δ* | *MATa his3Δ1 leu2Δ0 met15Δ0 ura3Δ0 gcn2Δ::KanMX4* | Open Biosystems |
| *hrk1Δ* | *MATa his3Δ1 leu2Δ0 met15Δ0 ura3Δ0 hrk1Δ::KanMX4* | Open Biosystems |
| *npr1Δ* | *MATa his3Δ1 leu2Δ0 met15Δ0 ura3Δ0 npr11Δ::KanMX4* | Open Biosystems |
| *psk1Δ* | *MATa his3Δ1 leu2Δ0 met15Δ0 ura3Δ0 psk1Δ::KanMX4* | Open Biosystems |
| *rck1Δ* | *MATa his3Δ1 leu2Δ0 met15Δ0 ura3Δ0 rck1Δ::KanMX4* | Open Biosystems |
| *rtk1Δ* | *MATa his3Δ1 leu2Δ0 met15Δ0 ura3Δ0 rtk11Δ::KanMX4* | Open Biosystems |
| *sak1Δ* | *MATa his3Δ1 leu2Δ0 met15Δ0 ura3Δ0 sak1Δ::KanMX4* | Open Biosystems |
| *ssn3Δ* | *MATa his3Δ1 leu2Δ0 met15Δ0 ura3Δ0 ssn3Δ::KanMX4* | Open Biosystems |
| *tos3Δ* | *MATa his3Δ1 leu2Δ0 met15Δ0 ura3Δ0 tos3Δ::KanMX4* | Open Biosystems |
| *ypk2Δ* | *MATa his3Δ1 leu2Δ0 met15Δ0 ura3Δ0 ypk2Δ::KanMX4* | Open Biosystems |
| y2203 | *MATa his3Δ1 leu2Δ0 met15Δ0 ura3Δ0 reg1Δ::KanMX4 Snf1-3xflag::HIS3* | In this study |
| y2211 | *MATa his3Δ1 leu2Δ0 met15Δ0 ura3Δ0 Tda1-TAP::HIS3* | Open Biosystems |
| y2230 | *MATa his3Δ1 leu2Δ0 met15Δ0 ura3Δ0 Tda1-3xflag::HIS3* | Open Biosystems |
| y2237 | *MATa his3Δ1 leu2Δ0 met15Δ0 ura3Δ0 snf1Δ::KanMX4 Tda1-3xflag::HIS3* | In this study |
| y2239 | *MATa his3Δ1 leu2Δ0 met15Δ0 ura3Δ0 cka1Δ::KanMX4 Tda1-3xflag::HIS3* | In this study |
| y2214 | *MATa his3Δ1 leu2Δ0 met15Δ0 ura3Δ0 Tda1-3xHA::KanMX6 Cka1-3xflag::HIS3* | In this study |
| y2215 | *MATa his3Δ1 leu2Δ0 met15Δ0 ura3Δ0 Tda1-3xHA::KanMX6 Snf1-3xflag::HIS3* | In this study |
| y2220 | *MATa his3Δ1 leu2Δ0 met15Δ0 ura3Δ0 Tda1-3xHA::KanMX6* | In this study |
| w303-1A | *MAT*a *leu2-3,112 trp1-1 can1-100 ura3-1 ade2-1 his3-11,15* | |
| yw001 | *MAT*a *leu2-3,112 trp1-1 can1-100 ura3-1 ade2-1 his3-11,15 tda1Δ::HIS3* | In this study |
| yw003 | *MAT*a *leu2-3,112 trp1-1 can1-100 ura3-1::[ADH1pr-Tda1-3xflag URA3] ade2-1 his3-11,15 tda1Δ::HIS3* | In this study |
| yw005 | *MAT*a *leu2-3,112 trp1-1 can1-100 ura3-1::[ADH1pr-Tda1 S483A/T484A-3xflag URA3] ade2-1 his3-11,15 tda1Δ::HIS3* | In this study |
| yw011 | *MAT*a *leu2-3,112 trp1-1 can1-100 ura3-1 ade2-1 his3-11,15 tda1Δ::HIS3 snf1Δ::KanMX6* | In this study |
| yw013 | *MAT*a *leu2-3,112 trp1-1 can1-100 ura3-1 ade2-1 his3-11,15 tda1Δ::HIS3 cka1Δ::KanMX6* | In this study |
| yw022 | *MAT*a *leu2-3,112 trp1-1 can1-100 ura3-1::[ADH1pr-Tda1-3xflag URA3] ade2-1 his3-11,15 tda1Δ::HIS3 cka1Δ::KanMX6* | In this study |
| yw018 | *MATa leu2-3,112 trp1-1 can1-100 ura3-1::[ADH1pr-Tda1-3xflag URA3] ade2-1 his3-11,15 tda1Δ::HIS3 snf1Δ::KanMX6* | In this study |
| yw014 | *MAT*a *leu2-3,112 trp1-1 can1-100 ura3-1::[ADH1pr-Tda1-3xflag-cMycNLS URA3] ade2-1 his3-11,15 tda1Δ::HIS3* | In this study |
| yw020 | *MAT*a *leu2-3,112 trp1-1 can1-100 ura3-1::[ADH1pr-Tda1-3xflag-cMycNLS URA3] ade2-1 his3-11,15 tda1Δ::HIS3 snf1Δ::KanMX4* | In this study |
| yw016 | *MAT*a *leu2-3,112 trp1-1 can1-100 ura3-1::[ADH1pr-Tda1-3xflag-cMycNLS URA3] ade2-1 his3-11,15 tda1Δ::HIS3 cka1Δ::KanMX6* | In this study |

**Supplemental Table S2. Yeast strains used in this study**
